# Role of an Atypical Cadherin Gene, *Cdh23* in Prepulse Inhibition and Implication of *CDH23* in Schizophrenia

**DOI:** 10.1101/2020.10.29.360180

**Authors:** Shabeesh Balan, Tetsuo Ohnishi, Akiko Watanabe, Hisako Ohba, Yoshimi Iwayama, Manabu Toyoshima, Tomonori Hara, Yasuko Hisano, Yuki Miyasaka, Tomoko Toyota, Chie Shimamoto-Mitsuyama, Motoko Maekawa, Shusuke Numata, Tetsuro Ohmori, Tomomi Shimogori, Yoshiaki Kikkawa, Takeshi Hayashi, Takeo Yoshikawa

## Abstract

We previously identified quantitative trait loci (QTL) for prepulse inhibition (PPI), an endophenotype of schizophrenia, on mouse chromosome 10 and reported *Fabp7* as a candidate gene from an analysis of F2 mice from inbred strains with high (C57BL/6N; B6) and low (C3H/HeN; C3H) PPI levels. Here, we reanalyzed the previously reported QTLs with increased marker density. The highest LOD score (26.66) peaked at a synonymous coding and splice-site variant, c.753G>A (rs257098870), in the *Cdh23* gene on chromosome 10; the c.753G (C3H) allele showed a PPI-lowering effect. Bayesian multiple QTL mapping also supported the same variant with a posterior probability of 1. Thus, we engineered the c.753G (C3H) allele into the B6 genetic background, which led to dampened PPI. We also revealed an e-QTL (expression-QTL) effect imparted by the c.753G>A variant for the *Cdh23* expression in the brain. In a human study, a homologous variant (c.753G>A; rs769896655) in *CDH23* showed a nominally significant enrichment in individuals with schizophrenia. We also identified multiple potentially deleterious *CDH23* variants in individuals with schizophrenia. Collectively, the present study reveals a PPI-regulating *Cdh23* variant and a possible contribution of *CDH23* to schizophrenia susceptibility.

## Introduction

Deciphering neurobiological correlates for behavioral phenotypes in terms of genetic predispositions that affect molecular, cellular and circuit-level functions is crucial for understanding the pathogenesis of psychiatric disorders.^1^ However, polygenicity and the inherent limitations of psychiatric diagnosis based on the subjective experiences of patients have impeded this endeavor.^2, 3^ With efforts to classify psychiatric disorders based on the dimensions of observable behavioral and neurobiological measures, endophenotype-based approaches for elucidating genetic liability have attracted interest because these approaches could mitigate clinical heterogeneity.^4, 5^

Among the behavioral endophenotypes, the prepulse inhibition (PPI) of acoustic startle response, a reflection of sensorimotor gating, has been consistently reported to be dampened in psychiatric disorders, particularly in schizophrenia.^6, 7^ As a robust and heritable endophenotype in schizophrenia,^8-11^ the genes/variants conferring the risk of dampened PPI can help to elucidate the genetic architecture of schizophrenia.^12, 13^ We have previously performed a large-scale quantitative trait loci (QTL) analysis for PPI and mapped six major loci through an analysis of 1,010 F2 mice derived by crossing selected inbred mouse strains with high (C57BL/6NCrlCrlj: B6Nj) and low (C3H/HeNCrlCrlj: C3HNj) PPI.^14^ Among these six loci, the chromosome-10 QTL showed the highest logarithm of odds (LOD) score, and we identified fatty acid-binding protein 7 (*Fabp7*) as one of the candidate genes in the PPI regulation and schizophrenia pathogenesis.^14, 15^ However, this high LOD score cannot be explained by a single gene, considering the low phenotypic variance attributed to the individual genes/markers, suggesting the presence of additional causative gene(s). ^16^ Therefore, in the current study, we aimed to (i) delineate additional gene(s) that regulate the PPI phenotype by reanalyzing the QTL with higher marker density, (ii) validate the candidate gene variant(s) by analyzing causal allele knock-in mice, and (iii) examine the potential role of candidate gene in schizophrenia susceptibility.

## Materials and Methods

### Animals

All animal experiments were approved by the Animal Ethics Committee at RIKEN. For the QTL analysis, we genotyped F2 mice from our previous study.^14^ For other experiments, C57BL/6NCrl (B6N) and C3H/HeNCrl (C3HN) mice were used (Japan’s Charles River Laboratories, Yokohama, Japan). Note that the previously used mouse strains C57BL/6NCrlCrlj (B6Nj) and C3H/HeNCrlCrlj (C3HNj) are no longer available because their supply ended in 2014.^16^

### Human DNA Samples

Studies involving human subjects were approved by the Human Ethics Committee at RIKEN. All participants in the genetic studies gave informed written consent. For resequencing the all protein-coding exons of the *CDH23* gene, a total of 1,200 individuals with schizophrenia (diagnosed according to DSM-IV) of Japanese descent (657 men, mean age 49.1±14.0 years; 543 women, mean age 51.1±14.4 years) were used. For subsequent genotyping, an additional 811 individuals with schizophrenia were included, for a total of 2,011 affected individuals (1,111 men, mean age 47.2 ± 14.1 years; 901 women, mean age 49.2 ± 14.7 years), along with 2,170 healthy controls (889 men, mean age 39.2 ±13.8 years; 1,281 women, mean age 44.6 ± 14.1 years). The samples were recruited from the Honshu region of Japan (the main island), where the population fall into a single genetic cluster.^17, 18^ Additionally, we also used genetic data of 8,380 healthy Japanese controls from the Integrative Japanese Genome Variation Database (iJGVD) by Tohoku Medical Megabank Organization (ToMMo) (https://ijgvd.megabank.tohoku.ac.jp/) to compare the allele frequencies of the identified variant.

### Human induced Pluripotent Stem Cells (hiPSCs)

To test the *CDH23* expression during neurodevelopment *in vitro*, hiPSC lines established from healthy controls (*n* = 4) were used.^19^ To analyze allele-specific expression of *CDH23* for the rs769896655 (c.753G>A) variant, another hiPSC line was established from a healthy individual (TKUR120) who was heterozygous for the variant. Establishment of hiPSCs and their differentiation to neurospheres and neurons were performed as described before.^20^

### QTL Analysis

Reanalysis of PPI-QTL was performed by increasing the density of the analyzed markers. To perform fine mapping, markers were selected To this end, exome sequencing of inbred C3HN and B6N mouse strains was performed, and single nucleotide variant (SNV) markers were selected from the six previously identified loci (chr1, chr3, chr7, chr10, chr11, chr13) for PPI (see supplementary methods).^14^ The 148 additionally selected SNV markers (supplementary table 1) were genotyped in 1,012 F2 mice by Illumina BeadArray genotyping (Illumina Golden Gate assay) on the BeadXpress platform as per the manufacturer’s instructions and were added to the analysis. QTL analysis was done by a composite interval mapping method^21^ using Windows QTL Cartographer v2.5 (https://brcwebportal.cos.ncsu.edu/qtlcart/WQTLCart.htm), and by Bayesian multiple QTL mapping (supplementary methods)

### Generation of Cdh23 c.753G Allele Knock-in Mice by CRISPR/Cas9n-mediated Genome Editing

The *Cdh23* c.753G allele knock-in mice in the B6N genetic background were generated by genome editing using the CRISPR/Cas9 nickase ^22-24^ as described in the supplementary methods and supplementary table 2.

### Analyses of PPI and Auditory Brainstem Response

The PPI test was performed according to previously published methods.^14, 15, 25^ Auditory brainstem response (ABR) was recorded as described previously (supplementary methods).^26^

### Cochlear Hair Cell Architecture Analysis

Scanning electron microscopy (SEM) of cochlear stereocilia was performed as described previously^27, 28^ in four ears from *Cdh23* c.753G allele knock-in mice and *Cdh23* c.753A, using a Hitachi S-4800 field emission scanning electron microscope at an accelerating voltage of 10 kV.

### Gene Expression Analysis

To detect the localization of *Cdh23* expression in the brain, we performed RNA *in situ* hybridization in mice (B6N and C3HN) and marmoset as described elsewhere (https://gene-atlas.brainminds.riken.jp/).^29, 30^ Expression of the target genes was measured by quantitative real-time RT-PCR (reverse-transcription polymerase chain reaction) or digital PCR using TaqMan Gene Expression Assays as described elsewhere (supplementary methods).^24, 31^ Splicing and exon skipping stemmed from the *Cdh23* c.753G>A variant were tested by semiquantitative RT-PCR normalized to *Gapdh* (supplementary table 3).

### Mutation Analysis of CDH23 in Schizophrenia by Targeted Next-generation Sequencing (NGS) using Molecular Inversion Probes (MIP)

Coding exons and the flanking exon-intron boundaries of the *CDH23* were sequenced using MIP as described previously (supplementary table 4).^32, 33^ Library preparation, sequencing and variant analysis were performed as described in the supplementary methods.

### Statistical Analysis

The data in the figures are represented as the mean ± SEM. Statistical analysis and graphical visualization was performed using GraphPad Prism 6 (GraphPad Software). The total sample size (*n*) and the analysis methods are described in the respective figure legends. Statistical significance was determined using a two-tailed Student’s *t*-test and the Holm-Sidak method. A *P* value of < .05 was considered statistically significant.

## Results

### Dense Mapping of QTL Reveals a High LOD Score on Chromosome 10 and Cdh23 as a Putative Candidate Gene

We reanalyzed the QTL for the PPI levels phenotyped in 1,010 F2 mice derived from F1 parents (B6NjC3HNj) from our previous study (supplementary figures 1A and 1B).^14^ To increase the density of the markers, in addition to the previously genotyped microsatellites, exome sequencing of C57BL/6NCrl (B6N) and C3H/HeNCrl (C3HN) mouse strains was performed (data not shown) and 148 strain-specific SNVs from the six previously reported loci on chr1, chr3, chr7, chr10, chr11 and chr13 were additionally genotyped (supplementary table 1).Composite interval mapping consistently revealed QTL signals with high LOD scores on chromosome 10 (supplementary figure 1C). The PPI at a prepulse level of 86 dB showed the highest LOD score, 26.66, which peaked at a synonymous coding and splice-site variant c.753G>A (rs257098870) in the cadherin 23 (*Cdh23*) gene (figures 1A and 1B, Supplementary Table 5). Additionally, Bayesian multiple QTL mapping also uncovered the same variant as the QTL with a posterior probability of 1 (figure 1C and supplementary table 6). Thus, we set out to pursue the role of *Cdh23* as a putative candidate gene.

**Fig. 1.**
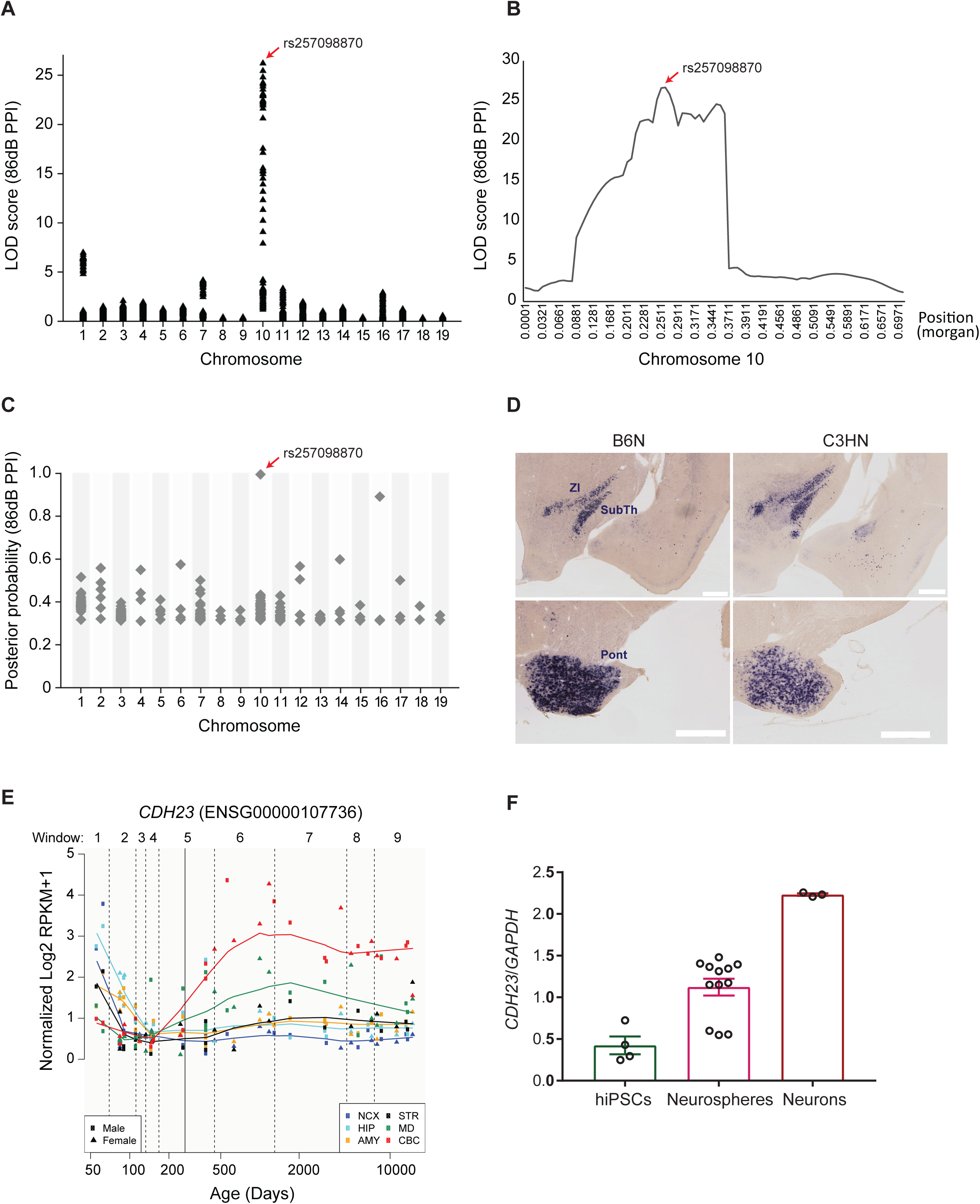
QTL mapping for PPI and expression analysis of *Cdh23* (A) LOD score calculated by composite interval mapping for prepulse level of 86 dB (B) Lod score distribution on chromosome 10. (C) Bayesian multiple QTL mapping for prepulse level of 86 dB. (D) RNA *in situ* hybridization in 3-week-old B6N and C3HN mice. ZI; zona incerta, SubTh; subthalamic nucleus, Pont; pontine regions (scale bar = 500 µm) (E) *CDH23* expression in humans (http://development.psychencode.org/) (F) *CDH23* expression in 4 hiPSC lines, neurospheres derived from each hiPSC line in triplicate, and neurons derived from one of the hiPSC lines in triplicate.

### Cdh23 is Predominantly Expressed in the Subthalamic and Pontine Regions of the Brain

Cdh23, an atypical cadherin, is a member of the cadherin superfamily, which encodes calcium-dependent cell-cell adhesion glycoproteins.^34, 35^ The role of *Cdh23* in cochlear hair cell architecture and hearing has been extensively studied.^36^ Although many members of the cadherin superfamily are expressed in neural cells and implicated in neurodevelopment, evidence for the role of *Cdh23* in neuronal functions is minimal. ^37 38, 39 40^ We evaluated *Cdh23* expression in the mouse brain by RNA *in situ* hybridization; the gene is predominantly expressed in subthalamic (including zona incerta and subthalamic nucleus) and pontine regions in both B6N and C3HN mouse strains (figure 1D) across the developmental stages (supplementary figure 2). Furthermore, a similar expression pattern was observed in the brain regions from the marmoset (*Callithrix jacchus*) (supplementary figure 3), indicating functional conservation across species. Previous reports have shown the potential role of these brain regions in sensorimotor gating.^41^ In humans, *CDH23* transcript expression was shown to be developmentally regulated across different brain regions, which was also observed in the *in vitro* neuronal differentiation lineage of hiPSCs (figures 1E and 1F).^19, 20^ These lines of evidence indicate potential roles for *Cdh23* in neuronal function and PPI.

### Cdh23 c.753G>A is the Only Distinct Coding Variant between B6N and C3HN Mouse Strains

Since a high LOD score was found for the SNV marker c.753G>A (rs257098870) in *Cdh23*, we reasoned that this variant or other flanking functional variants in the *Cdh23* gene might be responsible for the observed phenotypic effect. We resequenced all the coding and flanking intronic regions of the *Cdh23* gene in B6N and C3HN mice. However, no other variants were identified, except for the c.753G>A (rs257098870), on the 3’ end of exon 9 with the G allele in C3HN mice and the A allele in B6N mice (supplementary figures 4A-4D). The c.753A allele was previously known to cause age-related hearing loss (ARHL) by affecting cochlear stereocilia architecture ^28, 42, 43^. Although the variant does not cause an amino acid change (P251P), c.753A is known to elicit skipping of exon 9 in the inner ear.^42^ We also queried *Cdh23* genetic variations in C57BL/6NJ and C3H/HeJ mouse strains from Mouse Genomes Project (MGP) data (https://www.sanger.ac.uk/science/data/mouse-genomes-project/), which again showed c.753G>A as the only distinct coding variant between the strains (Supplementary Table 7). Moreover, inbred mouse strains with G allele identified from the MGP data (supplementary figure 4D) showed low PPI levels ^14^. Therefore, we reasoned that c.753G might contribute to the dampened PPI, and we set out to evaluate its role in PPI function.

### C3HN Strain-specific Cdh23 c.753G Allele Knock-in Mice Showed Reduced PPI Levels

To test the role of the *Cdh23* c.753G>A variant in modulating PPI, we generated a mouse model in the B6N genetic background in which the C3HN strain-specific *Cdh23* c.753G allele was knocked in using CRISPR/Cas9n-mediated genome engineering (supplementary figures 5A-5C). Mice homozygous for the knock-in allele (GG) and control littermates with the wild-type allele (AA) derived from heterozygous mice were used for the experiments. Since the A allele in the B6N strain results in the skipping of exon 9 and also causes ARHL, knock-in of the G allele in the B6 genetic background should reverse these events, and lead to dampening of the PPI.

We tested the splicing pattern of *Cdh23* exon 9 in the brain regions (subthalamic and pontine regions) where *Cdh23* was predominantly expressed, from knock-in mice homozygous for the G allele, control littermates homozygous for the A allele, and inbred B6N and C3HN mice. Exon 9 was almost completely retained in G allele-bearing mice, while the A allele elicited partial skipping of exon 9 (figure 2A). This was in line with the alternative splicing observed in inbred B6N and C3HN mouse brains (figure 2A). We found that the splicing pattern was tissue-specific, such that the expression of *Cdh23* transcript lacking exon 9 was lower in the pontine and subthalamic regions, than in the inner ear (supplementary figure 6). ^42^

**Fig. 2.**
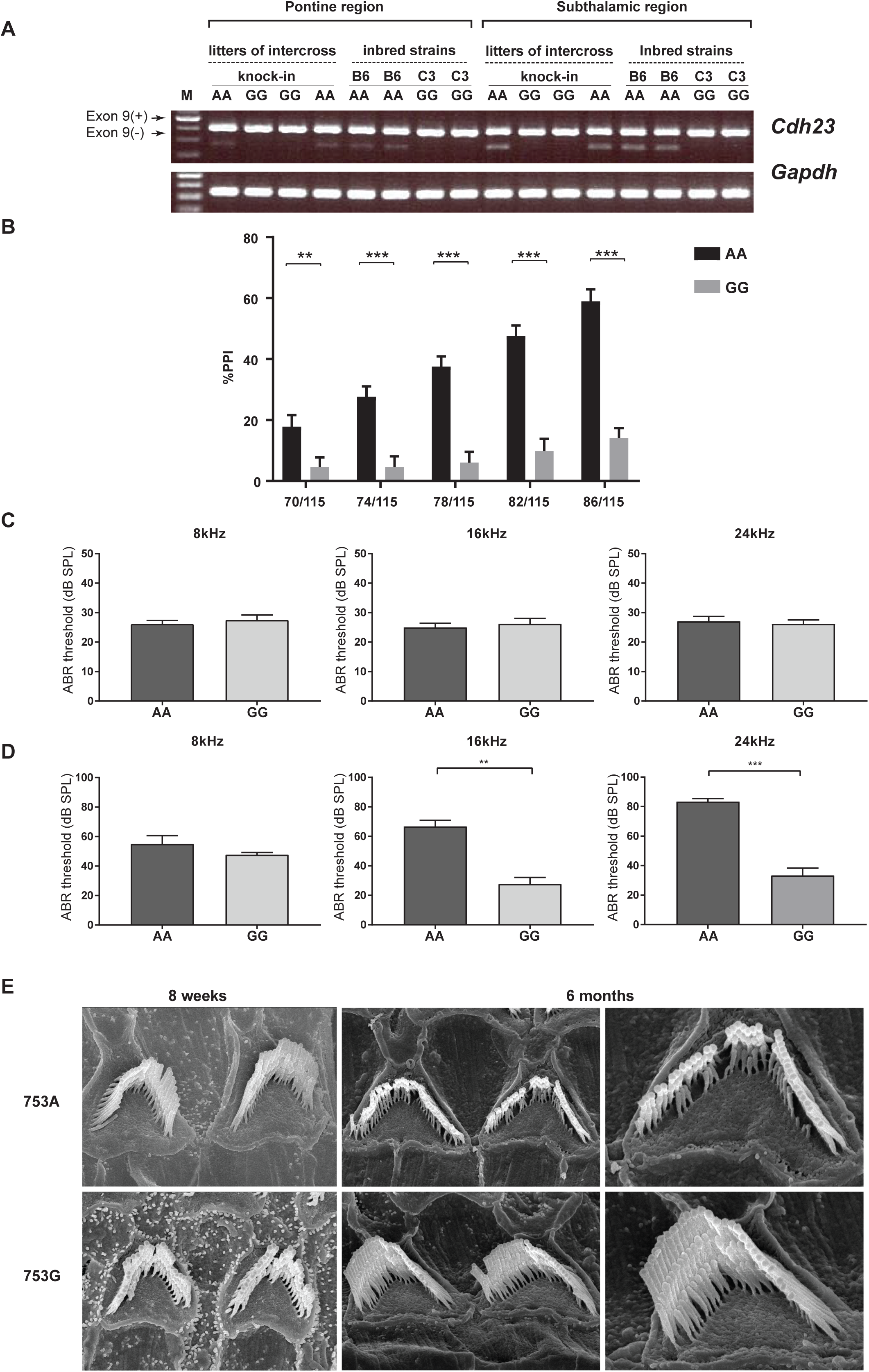
*Cdh23* c.753G allele knock-in in the B6N genetic background (A) Alternative splicing pattern. (B) PPI levels in *Cdh23* c.753G allele knock-in mice (*n* = 34) and c.753A allele littermates (*n* = 33) (13 weeks old). \*\**P* < .01, \*\*\**P* < .001 by Holm-Sidak method with alpha = 0.05 (C) ABR thresholds in *Cdh23* c.753AA (*n* = 16) and GG (*n* = 18) mice (13 weeks old). (D) ABR measurements in 1-year-old mice (AA; *n* = 6, GG; *n* = 9). \*\**P* < .01, \*\*\**P* < .001 by unpaired two-tailed Student’s *t*-test (E) Scanning electron microscope (SEM) images of cochlear hair cells.

Next, we tested the PPI levels in *Cdh23* c.753G allele knock-in mice and *Cdh23* c.753A allele litter mates (13 weeks old). The G allele knock-in mice showed lower PPI levels than the A allele mice in all the prepulse levels tested (figure 2B). Since the *Cdh23* c.753A allele is reported in ARHL, we further tested the hearing acuities of knock-in mice (GG) and control littermates (AA) to rule out any confounding effect. To this end, we examined the auditory brainstem response (ABR) thresholds for tone-pip stimuli at 8, 16 and 24 kHz in 13-week-old mice (the age at which PPI was measured) for both genotypes. We found no differences in the ABR threshold between AA and GG genotypes (figure 2C), thus supporting the conclusion that the observed impairment of PPI in the *Cdh23* c.753G allele knock-in mouse (or better PPI in the A allele) was not confounded by the deficits in hearing acuity, and that the *Cdh23* c.753A allele did not elicit hearing impairment at least up to 13 weeks of age. At 6 months of age, we observed a significantly higher ABR threshold in the *Cdh23* c.753A allele knock-in mouse than in the G allele, indicating the onset of ARHL in the AA genotype at this stage (figure 2D).^28, 43^ These observations were further corroborated with the evidence from scanning electron microscopy (SEM) images of cochlear hair cells (middle area), where the stereociliary architecture was intact in both genotypes at 8 weeks (figure 2E). The stereociliary morphology was preserved in 6-month-old *Cdh23* c.753G allele knock-in mice and the *Cdh23* c.753A allele littermates showed loss of hair cells indicating ARHL (figure 2E and supplementary figure 7). ^28, 44, 45^ These findings revealed that the C3HN strain-specific *Cdh23* c.753G is the causal allele that dampens PPI levels irrespective of hearing integrity.

### Potential Role of the Cdh23 c.753G>A Variant in Cdh23 Transcript Expression

To evaluate the functional role of the *Cdh23* c.753G>A variant, we examined the transcript expression levels of *Cdh23* in pontine and subthalamic regions using digital PCR. We observed significantly lower *Cdh23* transcript levels in the G allele carriers in both brain regions, indicating that c.753G>A acts as an expression QTL (e-QTL) (figure 3A). However, this trend was not observed in inbred adult B6N and C3HN mice (4-6 weeks-old) (figure 3B). Interestingly, differences in *Cdh23* transcript expression, between inbred B6N and C3HN mice, was observed during earlier stages of development particularly in E16.5 and P0 (figure 3C). Considering the developmentally regulated expression pattern of *Cdh23*, these results suggest that the e-QTL variant might act differentially during the early neurodevelopmental stages. Furthermore, it also points to the presence of epistatic interaction between the c.753 locus and genetic backgrounds for regulating the *Cdh23* transcript expression.

**Fig 3.**
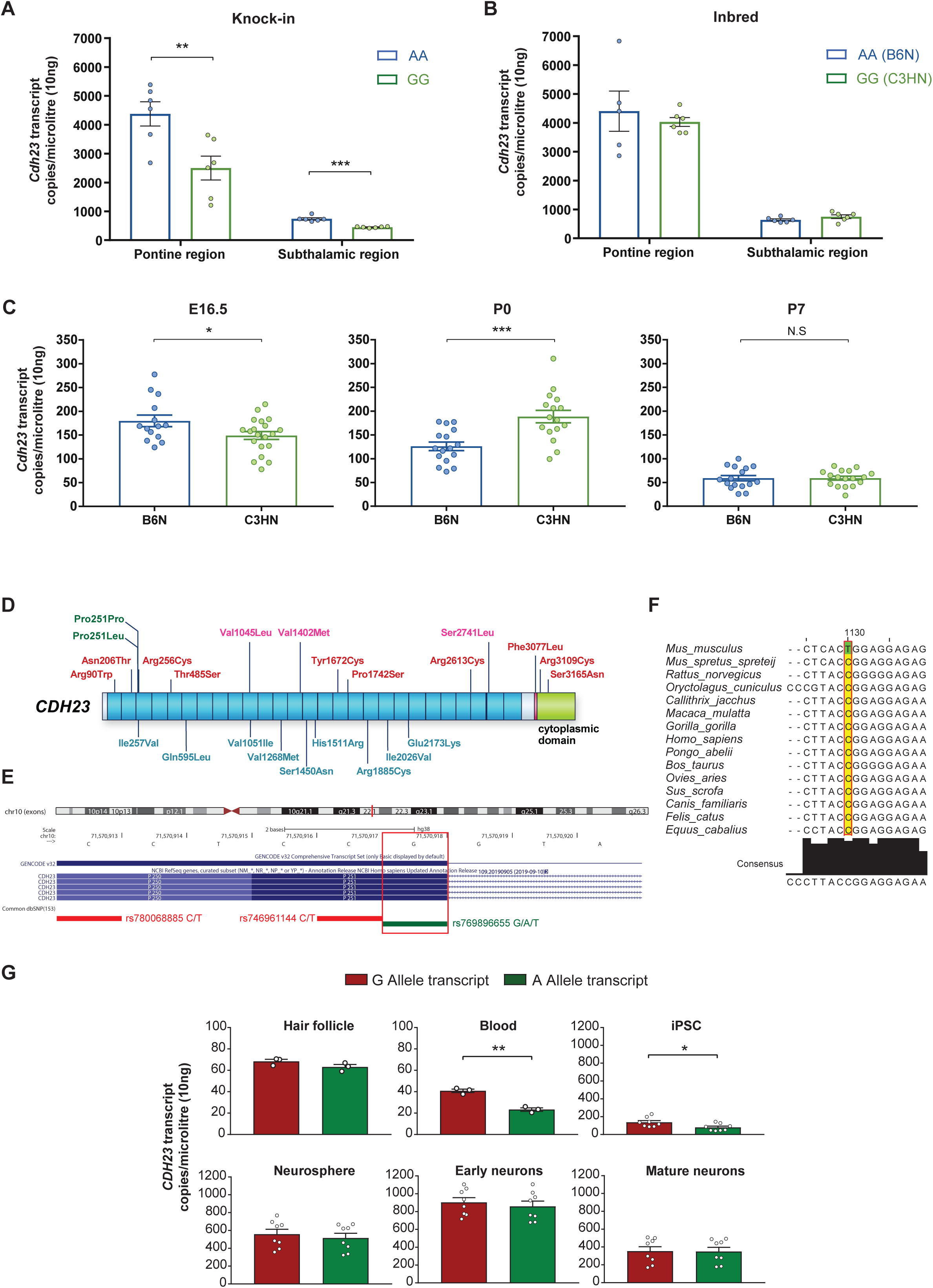
*Cdh23* expression and genetic variants in *CDH23* (A) Digital PCR-based *Cdh23* levels (*n* = 6 per group, age 4-6 weeks). \*\**P* < .01, \*\*\**P* < .001 by Holm-Sidak method with alpha =0.05 (B) Inbred B6N and C3HN mice (*n* = 6 per group, age 4-6 weeks). (C) *Cdh23* expression in whole cortex across the development (*n* = 14-17 per group). \**P* < .05, \*\*\**P* < .001 by unpaired two-tailed Student’s *t*-test (D) Novel rare variants in *CDH23* identified exclusively in schizophrenia cases (see supplementary figure 9). red; deleterious by ≥ 4 annotation tools, pink; deleterious by ≤ 3 tools, blue; not deleterious, green; homologous variant for *Cdh23* c.753G>A (rs769896655; c.753G>A/T P251P, and another variant resulting in P251L) (F) Allele conservation across species. (G) Allele-specific expression of *CDH23* in hair follicles (*n* = 1 in triplicate), peripheral blood samples (*n* = 1 in triplicate), hiPSCs (four lines, triplicate measurements), hiPSC-derived neurospheres, and neurons (early neurons; day 7, mature neuron; day 30 of differentiation) from a healthy subject heterozygous for the variant. \**P* < .05, \*\**P* < .01 by unpaired two-tailed Student’s *t*-test

### Role of CDH23 in Schizophrenia

Mounting evidence from genetic studies has indicated the potential role of *CDH23* in schizophrenia and other neuropsychiatric disorders.^46-50^ We therefore examined whether novel rare loss-of-function (LoF) variants in the coding regions of *CDH23* are enriched in Japanese schizophrenia cases, using molecular inversion probe (MIP)-based sequencing method (supplementary figure 8). Several patient-specific novel variants with potential deleterious effects predicted by *in silico* tools were identified in *CDH23*, although each variant was observed in 1-2 cases only (figure 3D and supplementary figure 9).

Interestingly, a homologous variant for *Cdh23* c.753G>A was observed in humans (rs769896655; c.753G>A; P251P), with G as the major allele (figures 3D-3F). The variant allele A was more frequent in the Japanese population than others in the gnomAD database (supplementary table 8).^51^ When we tested the enrichment of this variant in schizophrenia, a nominally significant over representation of A-allele in Japanese population was observed (schizophrenia vs. controls; *P* = .045) (table 1). The pooled analysis of schizophrenia samples [Japanese + Schizophrenia exome meta-analysis consortium, (SCHEMA); https://schema.broadinstitute.org/] and all controls, did not show any allelic association (*P* = .08) (supplementary table 9), which could be attributed to the rarity of variant allele A in other populations compared to the Japanese population.

**Table1.**
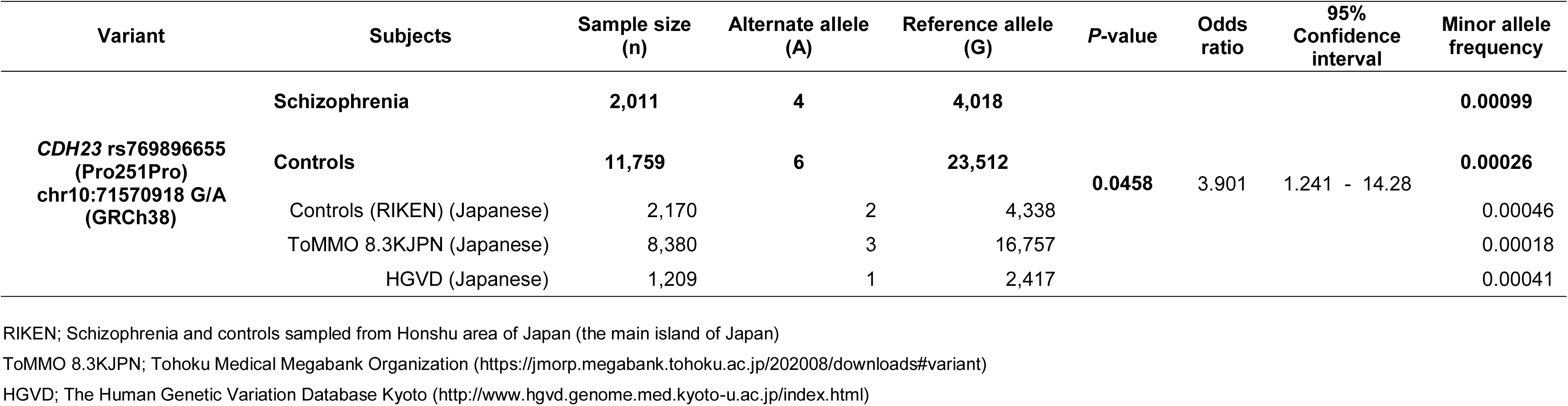
Association analysis of *CDH23* variant rs769896655 with schizophrenia

Allele-specific expression of the variant rs769896655 in the *CDH23* transcript was tested in hair follicles, peripheral blood samples, hiPSCs, hiPSC-derived neurospheres and neurons from a healthy subject who was heterozygous for the variant (supplementary figure 10A). A statistically significant preferential inclusion of the G allele was observed in *CDH23* transcripts from peripheral blood samples and hiPSCs (figure 3G), indicating that allele-specific expression is cell/tissue-specific and developmental stage-dependent.

## Discussion

By reanalyzing and fine-mapping mouse chromosome-10 PPI-QTL,^14^ we revealed *Cdh23* as a candidate for the PPI endophenotype. Our prior studies hinted at a possible role of *Cdh23*, but did not address this directly, as the gene is in the ARHL locus.^14, 16^ We showed here that the C3H/HeNCrl (C3HN) strain-specific G allele of the *Cdh23* c.753G>A variant is causal for low PPI, irrespective of hearing integrity. We also observed a potential e-QTL effect of the G allele in lowering *Cdh23* transcript expression in subthalamic and pontine regions in mice.

Several members of the cadherin superfamily govern multiple facets of neurodevelopment and function, but evidence for atypical cadherins is limited.^34^ The first evidence for the role of Cdh23 in neurodevelopment showed a lower number of parvalbumin-positive interneurons in the auditory cortex of *Cdh2*3-deficient mice, resulting from deficits in motility and/or polarity of migrating interneuron precursors during early embryonic development.^38^ Recently, *Cdh23* was also shown to be involved in mouse brain morphogenesis.^52^ Impaired PPI has also been attributed to neurodevelopmental deficits.^53^ Notably, the expression of *Cdh23* in the subthalamic and pontine regions is conserved across the species, as shown in mouse and marmoset in this study, and those regions are known to form the neural circuitry that regulates PPI.^41, 54-56^ Interestingly, deep brain stimulation (DBS) in the subthalamic region has been shown to improve PPI in rodent models and to ameliorate psychosis symptoms in individuals with Parkinson’s disease.^57, 58^

In this study, a homologous variant of mouse *Cdh23* c.753G>A in humans (rs769896655) showed a nominally significant enrichment, with overrepresentation of the A allele in Japanese population. The variant allele is rare in other populations, showing population specificity. Interestingly, a recent GWAS for the PPI phenotype in individuals with schizophrenia revealed a potential role of *CDH23*.^13^ Although not genome-wide significant, the lead variant in the *CDH23* interval is in considerable linkage disequilibrium with rs769896655 (supplementary figure 10B). It is also of note that schizophrenia-like psychosis was reported in Usher syndrome which is caused by mutations in *CDH23*.^48-50^

Although the c.753G allele was causal in lowering PPI in mice, the reason for the overrepresentation of A allele in schizophrenia cases is elusive. The wild mice sampled across different geographical locations showed the G allele in a homozygous state.^59^ It may be interesting to see whether low PPI is evolutionarily advantageous for mice to survive in the wild. A low PPI may help to enhance vigilance. The rarity of the variant A allele in humans, the lack of homozygotes and the population specificity indicate that the homologous variant might have originated recently. It is possible that *Cdh23*/*CDH23* may have a potential role in the PPI/schizophrenia phenotypes, but regulation of the phenotypes through the homologous variant might not be conserved. It is of note that the species specificity of variants in the manifestation of phenotypes is known in Parkinson’s disorder, where the disease-causing variant of alpha-synuclein in humans is commonly observed in rodents and other model organisms that do not display any pathological features.^60^ Regarding rs769896655 in *CDH23*, the rarity of variant allele A has limited the testing of genotype-phenotype correlation in humans.

In summary, the current study demonstrates the role of an atypical cadherin gene, *Cdh23*, in regulating PPI through a genetic variant without affecting hearing acuity, and a potential role of *CDH23* in schizophrenia. Large-scale studies to test the association of *CDH23* c.753G>A in schizophrenia cases are warranted in the Japanese population, where the variant allele is relatively prevalent.

## Supplementary Material

Supplementary material is available at *Schizophrenia Bulletin* online.

## Acknowledgements

We are grateful to the Animal Resources Development and Support Units for Bio-Material Analysis at RIKEN CBS Research Resources Division for animal maintenance, embryo manipulation, and DNA sequencing services.

