## Supplementary Information for "Role of an Atypical Cadherin Gene, *Cdh23* in Prepulse Inhibition and Implication of *CDH23* in Schizophrenia"

### Supplementary Methods

#### Exome sequencing in C3HN and B6N

Total DNA obtained from each one C3HN and B6N mouse was sheared into approximately 300-bp fragments using a Covaris sonicator (Covaris, Woburn, MA, USA). Paired-end exome libraries were prepared using the SureSelect Mouse All Exon Kit (Agilent) following the manufacturer's instructions. Sequencing was performed using the HiSeq 2000 platform (Illumina). Base calling was performed by the Illumina pipeline with default parameters. Obtained reads were mapped to the mouse genome (mm10) using the Burrows-Wheeler Aligner (BWA) version 0.5.9 with default parameters. Duplicate reads were marked with the Picard version 1.53 and excluded from downstream analyses. Coverage of the targeted bases was assessed by GATK DepthOfCoverage [1]. SNVs and indels were called by using the GATK UnifiedGenotyper3. The minimum phred-scaled confidence threshold was set as 50.

SNVs were selected from the regions ± 20 Mbp from the central position of each previously genotyped Sequence-Tagged Sites (STS) marker. One SNV marker from each megabase was primarily chosen using the following criteria: 1) specifically detected in either the B6N or C3HN strain, 2) homozygous, 3) with a PASS flag and 4) nearest to the central position of each megabase. If there were SNVs reported in dbSNP 128 within ± 0.25 Mbp of the central position of each megabase, we selected a marker from SNVs in dbSNP (that might not be the SNV nearest to the central position). The regions with no SNVs satisfying these criteria were skipped. If the distance between two markers was less than 0.25 Mbp, one marker was excluded. The list of the 148 selected SNV markers is shown in Supplementary Table 1.

**QTL analysis: Bayesian method**

Reanalysis of PPI-QTL was performed by increasing the density of the analyzed markers. To perform fine mapping, single nucleotide variant (SNV) markers were selected from the six previously identified loci (chr1, chr3, chr7, chr10, chr11, chr13) for PPI [2]. To this end, exome sequencing of inbred C3HN and B6N mouse strains was performed and markers were selected as described in the supplementary methods. The 148 additionally selected SNV markers (Supplementary Table 1) were genotyped in 1,012 F2 mice by Illumina BeadArray genotyping (Illumina Golden Gate assay) on the BeadXpress platform as per the manufacturer's instructions and were added to the analysis. First, we applied a composite interval mapping method [3] for 86 dB-PPI to scan relevant QTLs using Windows QTL Cartographer v2.5 (https://brcwebportal.cos.ncsu.edu/qtlcart/WQTLCart.htm) on autosomal chromosomes only, as no QTL signal was detected on the X chromosome in the previous analyses [2]. In the present analysis, the gender of the mice was taken into consideration. The results were expressed as LOD scores, and a threshold of LOD score corresponding to a genome-wide significance level of 1% was determined to be 4.16 with 1000 repetitions of the permutation test. Subsequently, Bayesian multiple QTL mapping was conducted for fine mapping of QTLs at the 86 dB prepulse level. In Bayesian method, we located a putative QTL on each marker and estimated the posterior probability that the QTL had a nonzero significant effect. The phenotype of the $i^{th}$ F2 mouse, $y_i$, was written as the following statistical model:

$$y_i = \mu + s_i b + \sum_{l=1}^{N} (\gamma_l u_{il} a_l + \eta_l v_{il} d_l) + e_i$$

where $\mu$ was the intercept of the model, $b$ was the gender effect, $s_i$ was a covariate indicating the gender of the $i$th mouse with $s_i$ =0 and 1 corresponding to male and female, respectively,

$N$ was the number of markers on which putative QTLs were located, $u_{il}$ and $v_{il}$ were the covariates indicating the genotype of the $i$th mouse at the $l$th marker, with $u_{il}$ =1, 0, -1 and $v_{il}$ = 0, 1, 0 for homozygous for the B6 type and heterozygous and alternative homozygous for the C3 type, respectively, $a_l$ and $d_l$ were additive and dominance effects of the $l$th marker, $\gamma_l$ and $\eta_l$ were indicator variables with $\gamma_l$ = 1 or 0 meaning the inclusion or exclusion of the additive effect at the QTL in the model fitting, respectively, and analogously, $\eta_l$ =1 or 0 for inclusion or exclusion of dominance effect, and $e_i$ was the residual error assumed to follow a normal distribution with mean 0 and variance $\sigma_e^2$.

In a Bayesian framework, these parameters and indicator variables were evaluated based on their posterior distributions constructed from the prior distributions and the distribution of observed samples, including $y_i$, $u_{il}$ and $v_{il}$ ($i$=1,2,…,$n$; $l$=1,2,…,$N$). The posterior distributions can be empirically established by sampling values of the parameters and indicator variables via the Markov chain Monte Carlo (MCMC) sampling procedure, which is time-consuming. Instead, we applied a variational approximation method to effectively obtain the posterior expectations of the parameters and indicator variables [4]. This variational Bayes method was originally utilized in the context of genomic selection [5], where a model predicting breeding values from SNP genotypes, which is analogous to the model considered here, is constructed to enable individuals with superior genetic performance to be effectively selected based on their SNP genotypes in animal and plant breeding [4]. The existence of QTLs with significant effects was judged using the posterior expectations of indicator variables, $\gamma_l$ and $\eta_l$, for additive and dominance effects, which were regarded as the posterior probabilities of the putative QTLs being fitted in the model. As a result of this variational Bayes analysis, we found no QTLs with a significant dominance effect; thus, only the plot of the posterior expectations of $\gamma_l$ is shown in Figure 1C.

**Conservation analysis of *Cdh23* c.753 G>A variant**

Since the only coding variant that differed between the B6N and C3H mouse in the Chromosome 10 QTL region was *Cdh23* c.753 G>A; (rs257098870; chr10:60530947-60530947; mm10/GRCm38), we further compared the conservation of this variant/region among 15 eutherian mammals (*Mus musculus, Mus Spretus* (SPRET/Eij), *Rattus norvegicus, Oryctolagus cuniculus, Callithrix jacchus, Macaca mulatta, Gorilla gorilla, Homo sapiens, Pongo abelii, Bos taurus, Ovis aries, Sus scrofa, Canis familiaris, Felis catus, Equus caballus*) and rodents; which included 15 inbred mouse strains (C57BL/6NJ, NZO/HlLtJ, A/J, BALB/cJ, AKR/J, C3H/HeJ, CBA/J, DBA/2J, FVB/NJ, NOD/ShiLtJ, 129S1/SvImJ, LP/J, WSB/EiJ, CAST/EiJ, and PWK/PhJ), along with *Mus Spretus and Rattus norvegicus*. Alignments for the variant and flanking regions were downloaded from Ensembl (https://www.ensembl.org/ ) and visualized using Jalview (https://www.jalview.org/).

**Generation of *Cdh23* c.753G allele knock-in mice by CRISPR/Cas9n-mediated genome editing**

Briefly, B6N zygotes obtained by *in vitro* fertilization were microinjected with the cocktail constituting 5 ng/mL Cas9 nickase mRNA (System Biosciences, Mountain View, CA), 5 ng/mL each of two short guide RNAs (sgRNAs) (Supplementary Table 2), which were synthesized *in vitro* (T7 gRNA Smart Nuclease Synthesis Kit, System Biosciences) according to the manufacturer's instructions, and 5 ng/mL single-stranded oligodeoxynucleotide (ssODN) (Supplementary Table 2). Injected zygotes were transplanted into the uteruses of pseudopregnant dams, and the resulting pups were obtained by cesarean section. The target region of the *Cdh23* gene was directly sequenced from the PCR products amplified from the template DNAs extracted from the tails of the pups. Additionally, the PCR products were subcloned and sequenced in founders with knocked-in alleles. Promising founders were selected, and upon their reaching sexual maturity, *in vitro* fertilization was performed with B6N strain-derived oocytes to check germline transmission and to obtain mice with heterozygous mutated alleles. Furthermore, the heterozygous mice were intercrossed to produce homozygote knock-in and wild-type control littermates for further experiments. Routine genotyping by Sanger sequencing was performed to verify the allele knock-in.

**Analysis of auditory brainstem response**

Briefly, mice were anesthetized by intraperitoneal injection of sodium pentobarbital (Somunopentil; 60-80 mg/kg) diluted in saline and were placed on a heating pad to maintain body temperature at 37°C during ABR evaluation. Needle electrodes were placed subcutaneously into the vertex (reference), right ear (active), and left ear (ground). The tone stimulus was produced by a speaker (ES1 spc; BioResearch Center Nagoya, Japan) probe inserted into the auditory canal of the right ear. The ABR thresholds from right ears in 13-week-old and 6-month-old mice (*Cdh23* c.753G>A; AA vs GG genotypes) were measured with tone-pip stimuli at 8, 16 and 24 kHz generated by System3 (TDT). The resulting ABR waveforms were bandpass-filtered (< 3 kHz and > 100 kHz), amplified 1,000 times (AC PreAmplifier, P-55, Astro-Med Inc.) and recorded using the PowerLab 2/25 System (AD Instruments) for 10 microseconds. A total of 500 recordings were averaged. The waveform data were analyzed by LabChart v.7 (AD Instruments). The ABR thresholds were obtained for each stimulus by reducing the sound pressure level (SPL) for 10 dB steps (80 dB to 10 dB) to identify the lowest level at which an ABR pattern could be reliably detected. The ABR threshold was confirmed for consensus by independent evaluation of three investigators blinded to the genotype of the tested mice.

**Gene expression analysis**

Total RNA was extracted using the miRNeasy Mini kit (QIAGEN GmbH, Hilden, Germany), and single-stranded cDNA was synthesized using the SuperScript VILO cDNA synthesis kit according to the manufacturer's instructions. The mRNA levels for the target genes (Supplementary Table 3) were quantified by real-time quantitative RT-PCR using TaqMan Gene Expression Assays, performed in triplicate, based on the standard curve method, normalized to GAPDH/Gapdh as an internal control. To test the role of the *Cdh23* c.753G>A variant as a cis expression QTL (e-QTL) for *Cdh23* expression, digital PCR was performed in mouse brain samples from the subthalamic region/zona incerta and pontine region using standard procedures and TaqMan Gene Expression Assays in a QuantStudio™ 3D Digital PCR System (Life Technologies Co., Carlsbad, CA, USA). Values outside the mean ± 2 SD in the group were considered outliers and were omitted from the analyses.

**Design, library preparation, sequencing for MIP**

Genomic DNA was isolated from blood samples obtained from human subjects using standard methods. MIPs were designed for the *CDH23* gene using MIPgen [6] (http://shendurelab.github.io/MIPGEN/), targeting the coding exons and the flanking exon-intron boundaries of the gene (GRCh37 build) and covering all transcripts. A total of 152 MIPs were designed for *CDH23*, aligning to the design parameters as reported previously [7].

A total 50 ng of genomic DNA, 250 fmol of 5'-phosphorylated MIPs mixture, 1.5 µL of 10X Ampligase DNA ligase buffer (Epicentre), 0.3 µM dNTP (Invitrogen, Carlsbad, CA, USA), 0.2 µL Hemo Klentaq (New England Biolabs, NEB, Ipswich, MA, USA) and 1 U Ampligase (Epicentre biotechnologies, Madison, WI, USA) were added to molecular biology-grade water for a total of 15 µL. After denaturation (95°C) for 10 minutes and incubation for (60 °C) for 22 hours, linear probes and the remaining genomic DNA were removed by exonuclease treatment. Next, the captured material was amplified by PCR using barcoded reverse primers. At first, 48 samples PCR product were pooled, purified and sequenced by Illumina MiSeq (Illumina, San Diego, CA, USA) system with 150 bp paired-end reads. Depend on the read number of each MIPs, rebalanced the mixture ratio. After that, operated same method using rebalanced MIP-mixture, 384 samples (3 sets) PCR product were pooled, purified and sequenced by Illumina HiSeq2000 (Illumina) Rapid mode with 150 bp paired-end reads.

**Variant annotation and analysis**

In accordance with the Genome Analysis ToolKit (GATK) best practices [8], raw reads were mapped to the genome (GRCh37 build) using Burrows-Wheeler Aligner (BWA MEM) (0.7.5a-r405) which was further converted to .bam files using Picard-tools (1.137). The GATK (3.4-46) was used for indel realignment, base-quality recalibration, and variant discovery. Individual variant calling was performed with the GATK Haplotype Caller, followed by multi-sample genotyping, and subsequently the variants were hard filtered. Variants with Quality score (QUAL) < 30.0, quality by depth (QD) < 5.0, genotype quality (GQ) < 20.0, heterozygous allele balance (ABHet) > 0.75, homopolymer run (HRun) >= 4, and clustered variants were excluded from the analysis. Variants were further annotated using ANNOVAR for deleteriousness [9] and was filtered for only missense and loss of function (LoF) variants (stop gained, stop lost, start lost, and variants on splice donor and acceptor sites). In addition we also annotated the variants with REVEL tool [10]. Since we were interested in the variants exclusively present in cases, we excluded the variants that were present in Exome Aggregation Consortium (ExAC) (http://exac.broadinstitute.org/) for global population, Integrative Japanese Genome Variation Database (iJGVD) (3.5KJP) by Tohoku Medical Megabank Organization (ToMMo) (https://ijgvd.megabank.tohoku.ac.jp/) and Human Genetic Variation Database (HGVD) (http://www.hgvd.genome.med.kyoto-u.ac.jp/)

for Japanese population. The identified variants were further validated by the Sanger sequencing.

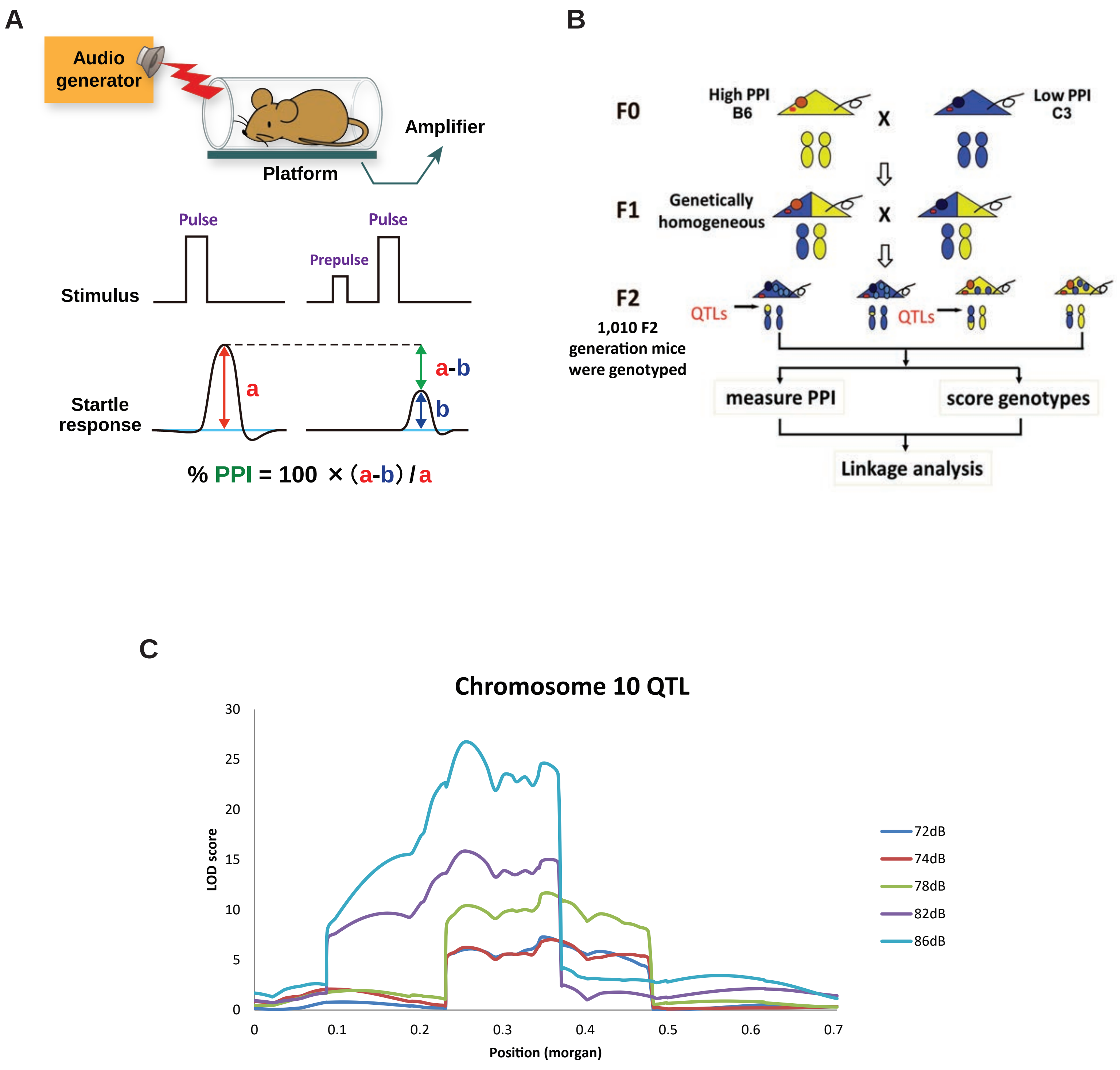


**Supplementary Figure 1: Experimental descriptions** (**A**) Schematic representation describing evaluation of prepulse inhibition (PPI) (**B**) linkage analysis scheme for quantitative trait loci (QTL) analysis (**C**) chromosome 10 QTL for PPI at different prepulse levels

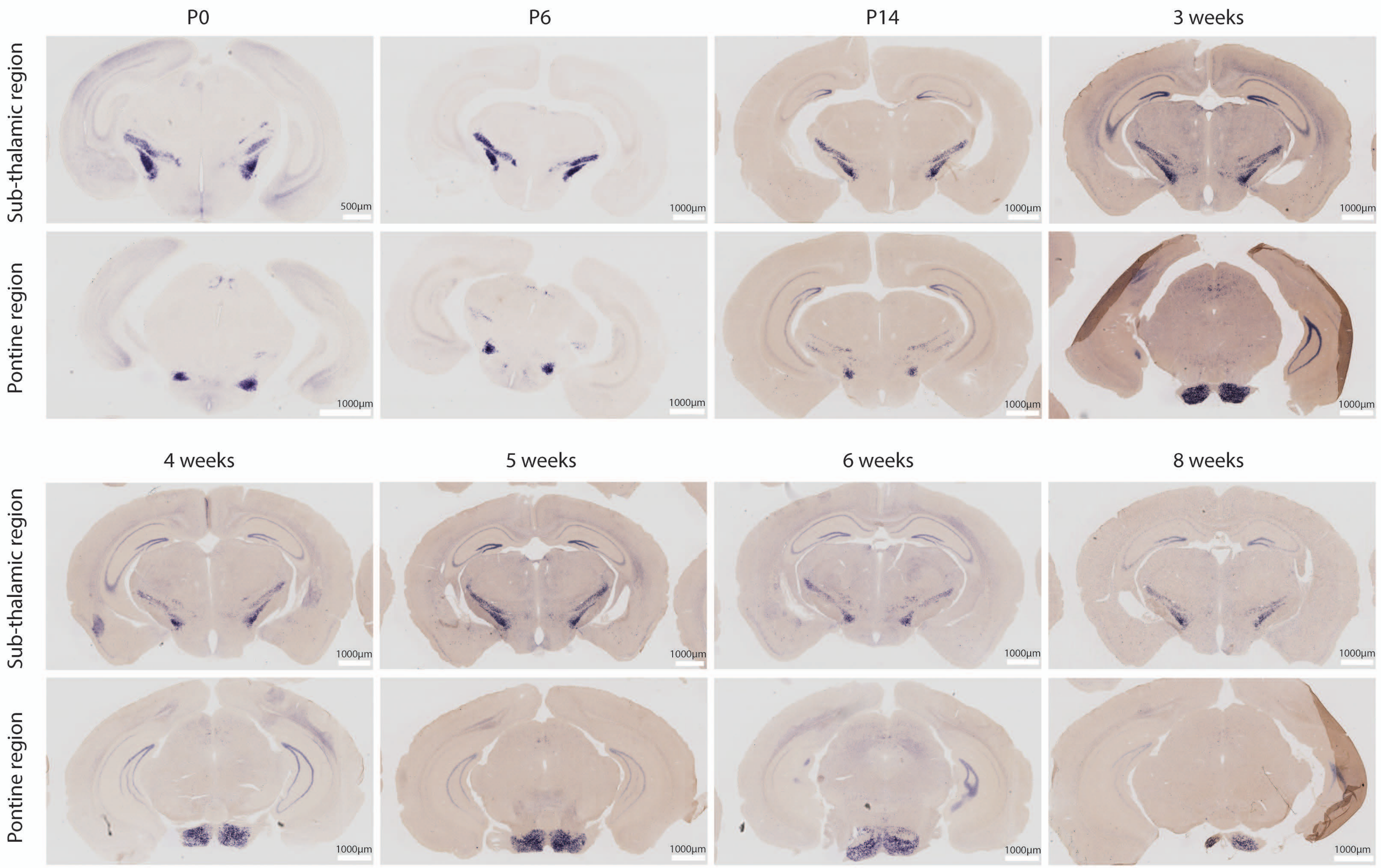


**Supplementary Figure 2: *Cdh23* expression analysis in mouse brain sections across development.** RNA *in situ* hybridization for *Cdh23* in B6N mouse showed expression in the brain particularly in subthalamic and pontine regions.

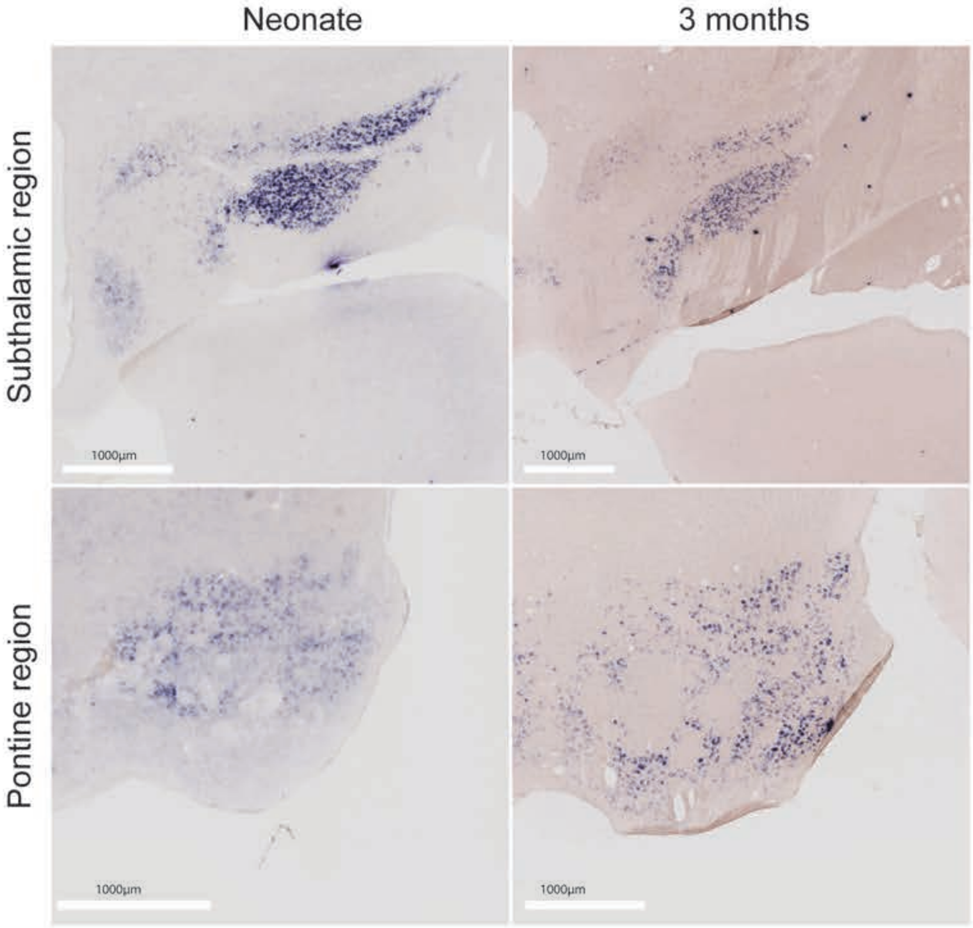


**Supplementary Figure 3: *Cdh23* expression in marmoset brain sections. (A)** RNA *in situ* hybridization data shows pattern of *Cdh23* expression is conserved in primates as evidenced from neonatal and 3 months marmoset (*Callithrix jacchus*) brain sections (https://gene-atlas.brainminds.riken.jp/)

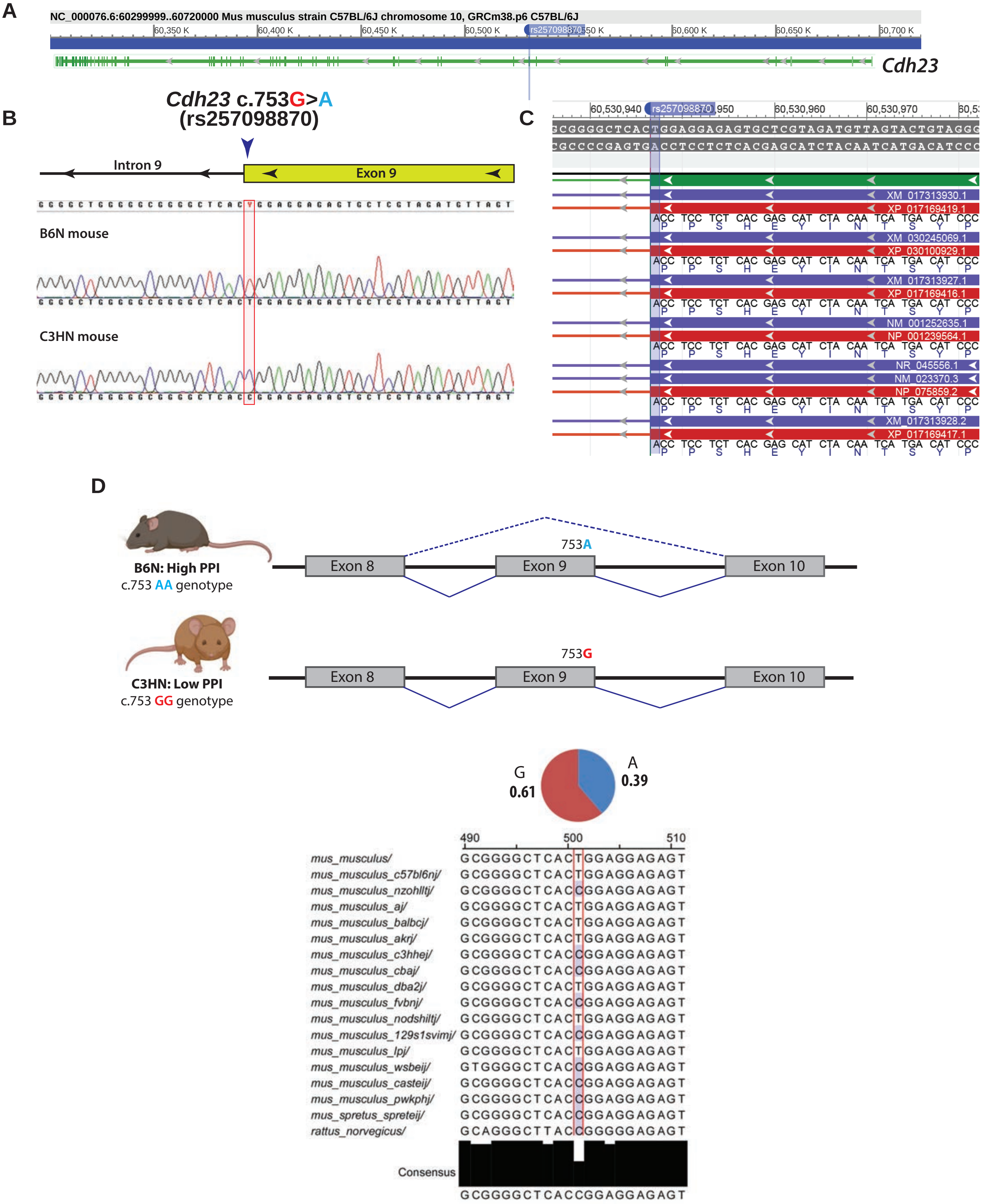


**Supplementary Figure 4: *Cdh23* gene and the putative causal variant c.753G>A (rs257098870).** **(A)** Gene structure of *Cdh23* gene with the putative causal variant marked **(B)** Electropherogram showing the c.753G>A variant alleles in B6N and C3HN mice strains **(C)** The variant (synonymous coding) does not cause amino acid change (P251P), **(D)** but results in tissue-dependent skipping of the exon 9, in varying levels. The frequency of the variant allele is relatively high as observed from the mouse genome project data, indicating strain specificity and lack of conservation

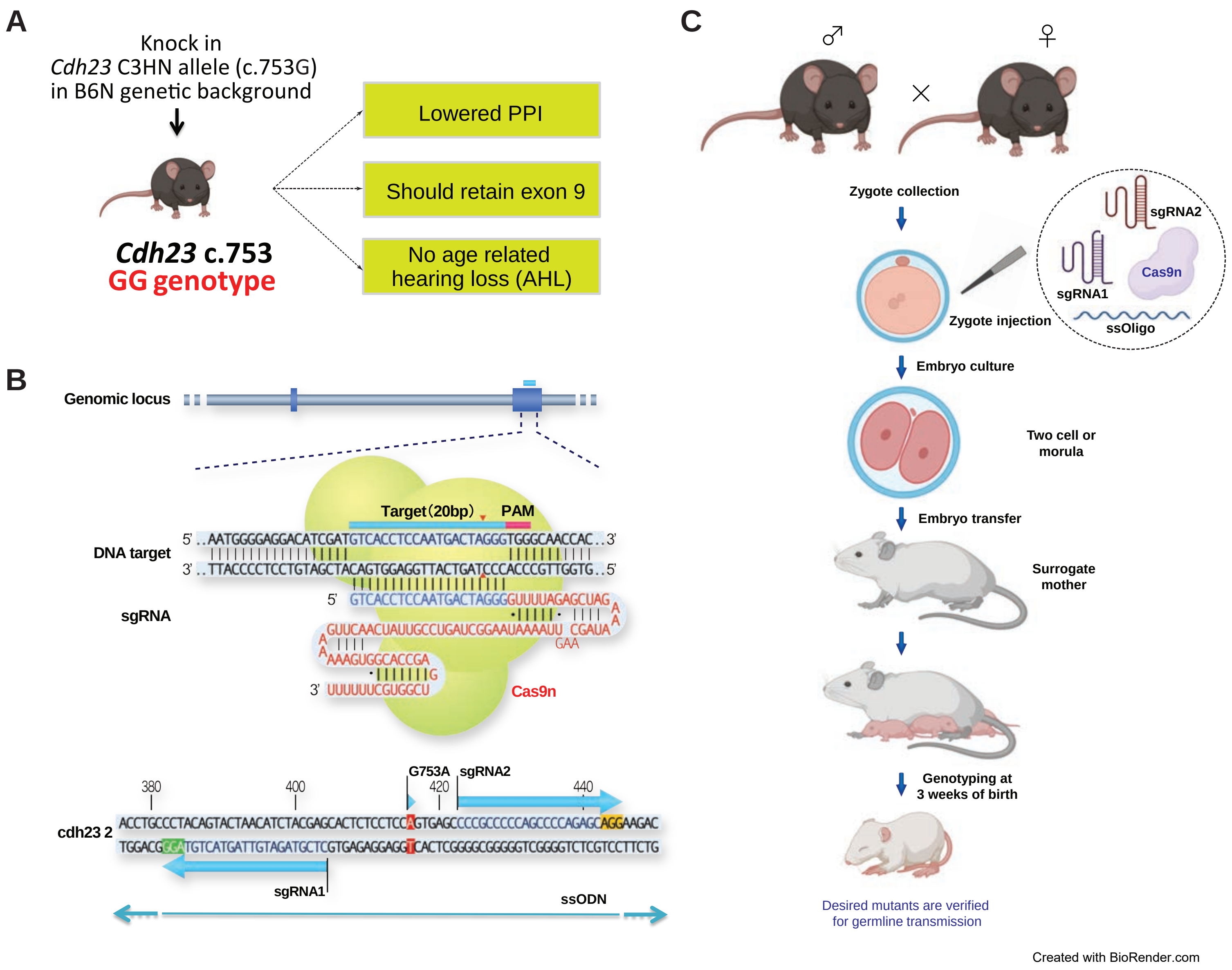


**Supplementary Figure 5: Experimental descriptions** (**A**) Schematic representation describing CRISPR-Cas9n mediated genomic editing for knocking in C3HN-specific G allele of *Cdh23* c.753G>A variant in B6N genetic background and the expected outcome of experiments by knocking in G allele. (**B**) Targeted genomic loci for genomic editing and designs for short guide RNA (sgRNA) and single-stranded oligodeoxynucleotide (ssODN) (**C**) Experimental strategy for generating genome edited mouse.

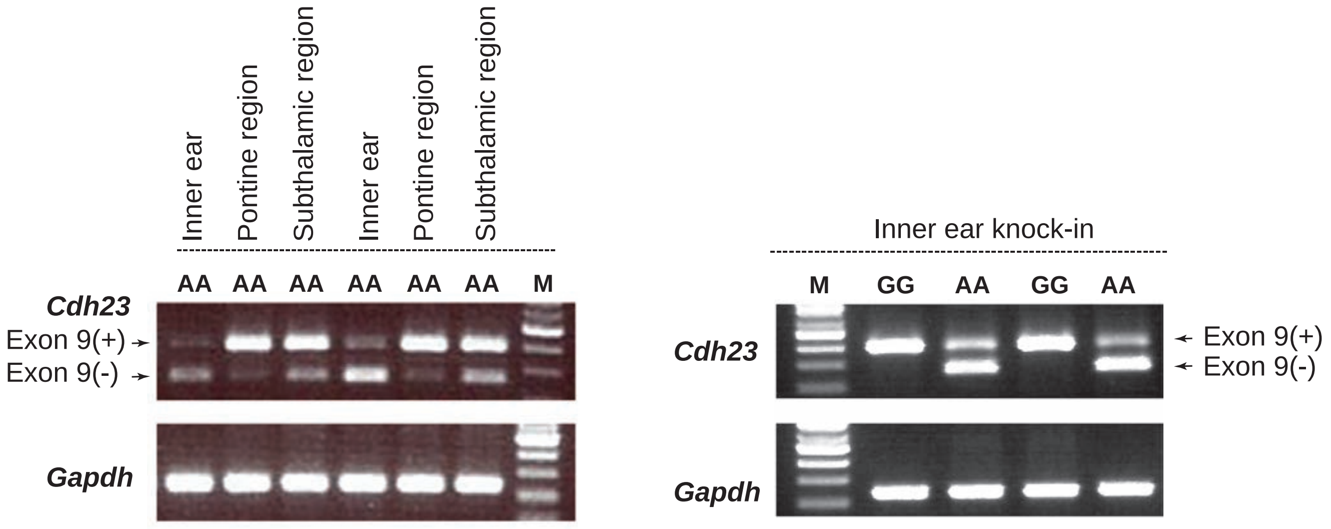


**Supplementary Figure 6: *Cdh23* alternative splicing in inbred B6N mouse strain and prepulse inhibition (PPI) levels in *Cdh23* c.753G allele knock-in mice.** Comparative analysis of *Cdh23* alternative splicing in the brain and inner ear from inbred B6N mouse strains showed tissue-specific alternative splicing of exon 9, which is lesser in brain (pontine and subthalamic region compared to the inner ear. The higher level of exon 9 containing transcripts in the inner ear from the *Cdh23* c.753G allele knock-in mouse was in agreement with the inbred strains with G allele, thus underscoring successful knock-in.

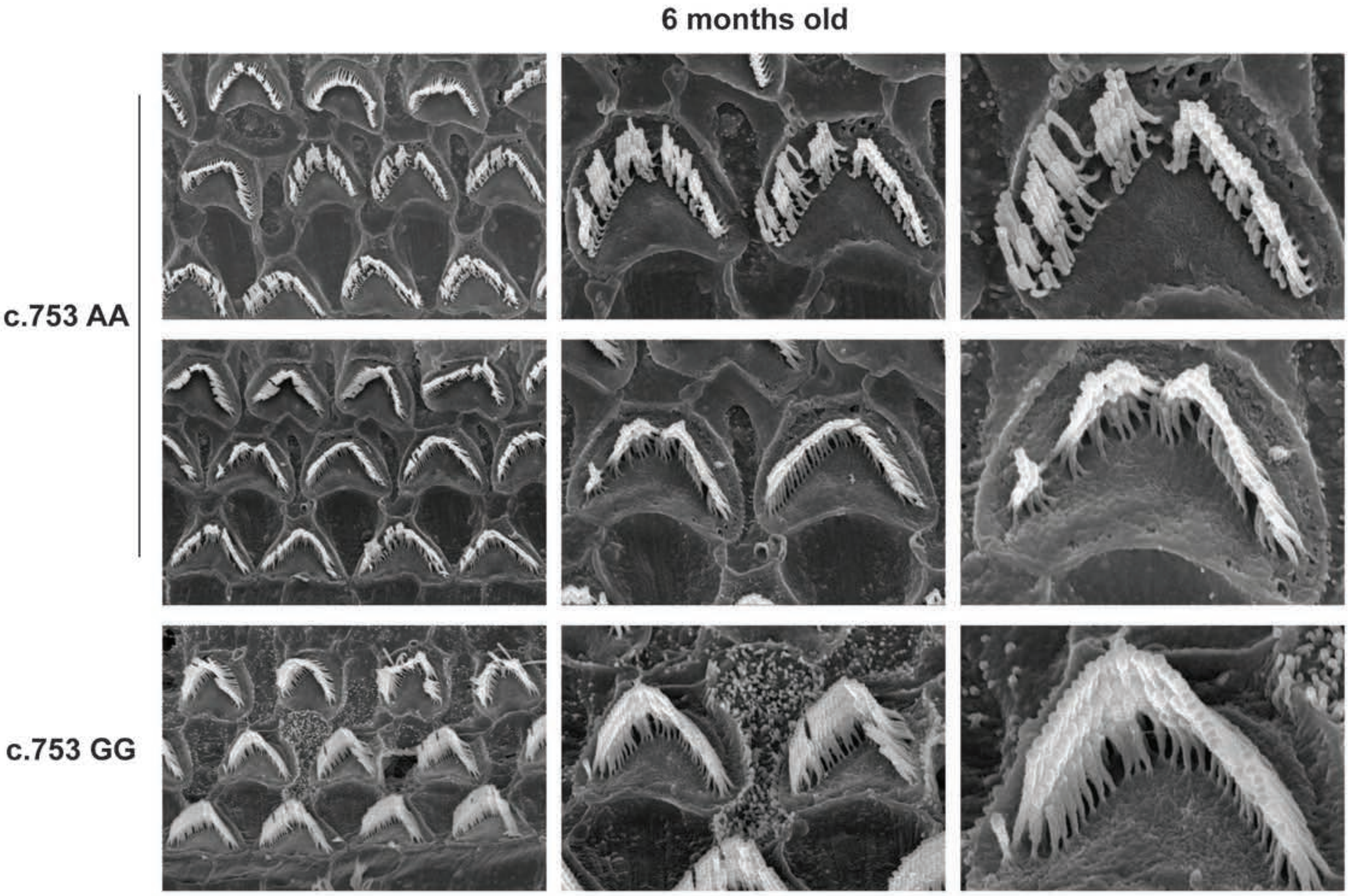


**Supplementary Figure 7: Scanning electron microscopy (SEM) images of hair cells.** Stereocillia morphology (middle area) is preserved in 6 months old *Cdh23* c.753G allele knock-in mouse (GG) in B6 genetic background, when compared to the *Cdh23* c.753A allele littermates (AA), which showed loss of hair cells, indicating age-related hearing loss

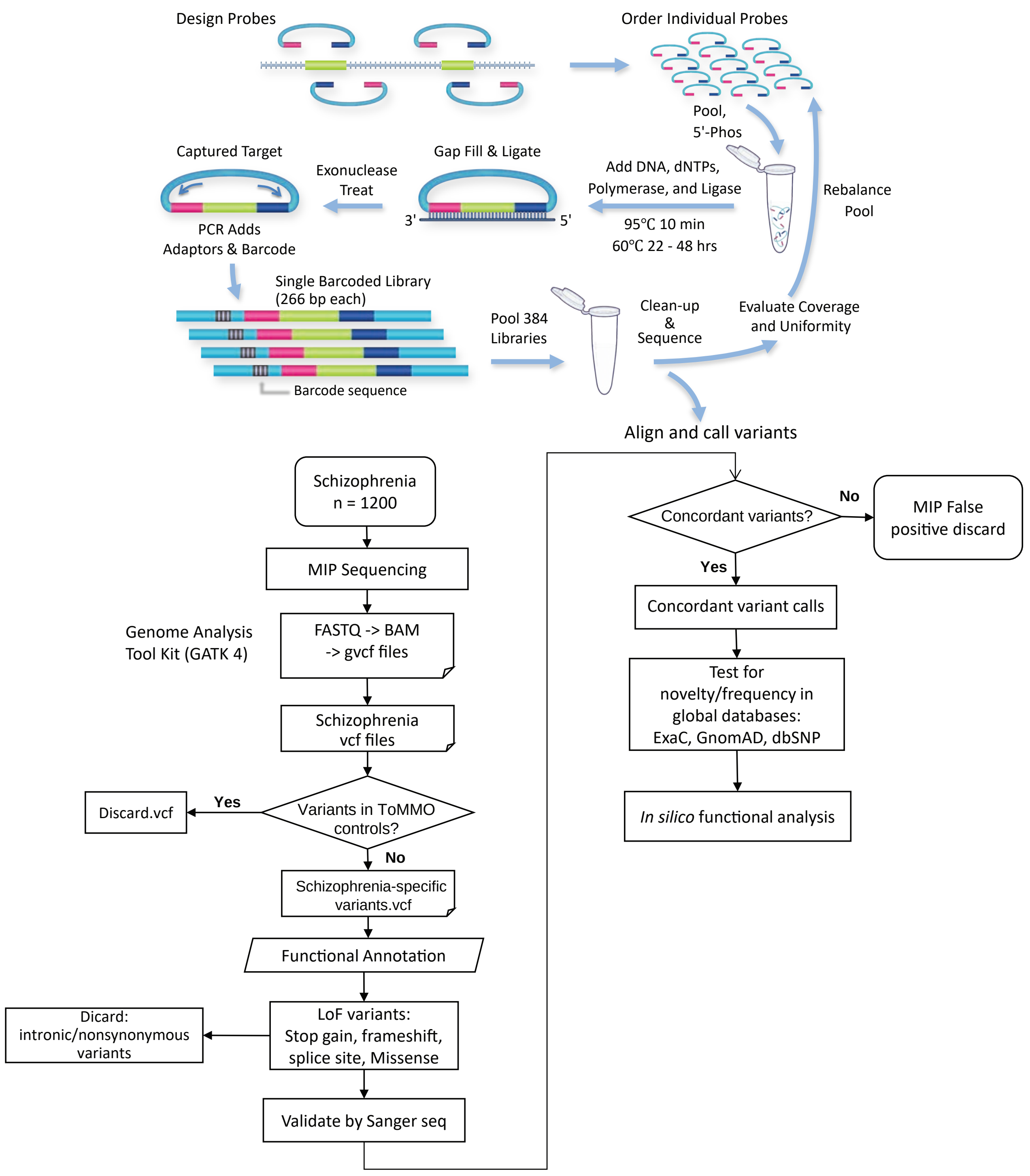


**Supplementary Figure 8: Role of *CDH23* in schizophrenia.** We tested whether novel rare LoF variants are present in Japanese schizophrenia cases, using molecular inversion probe (MIP) based sequencing as outlined.

#### *CDH23*: Cadherin related 23

| Chr | hg19_position | Ref | Alt | name | Gene | Annotations | | | | | | | | | REVEL_Scores | CADD_phred | GERP++_RS | Count_SCZ |
|---|---|---|---|---|---|---|---|---|---|---|---|---|---|---|---|---|---|---|
| 10 | 73537606 | A | G | p.Tyr1672Cys | CDH23 | D | D | D | D | D | H | . | D | D | 0.94 | 22.4 | 5.76 | 1 |
| 10 | 73539060 | C | T | p.Pro1742Ser | CDH23 | D | D | D | D | D | H | . | D | D | 0.741 | 28.8 | 4.88 | 1 |
| 10 | 73572350 | G | A | p.Ser3165Asn | CDH23 | D | D | D | D | D | L | D | D | D | 0.463 | 33 | 5.44 | 1 |
| 10 | 73269961 | C | T | p.Arg90Trp | CDH23 | D | D | D | D | D | M | T | T | T | 0.378 | 18.71 | 1.76 | 1 |
| 10 | 73563142 | C | T | p.Arg2613Cys | CDH23 | D | D | P | D | D | M | T | T | T | 0.486 | 16.16 | 4.23 | 1 |
| 10 | 73326686 | A | C | p.Asn206Thr | CDH23 | T | D | D | D | D | N | T | T | T | 0.246 | 18.21 | 5.6 | 1 |
| 10 | 73337683 | C | T | p.Arg256Cys | CDH23 | T | D | D | D | D | N | T | T | T | 0.249 | 20.7 | 5.79 | 2 |
| 10 | 73434873 | C | G | p.Thr485Ser | CDH23 | T | D | D | D | D | L | . | T | T | 0.172 | 16.85 | 5.72 | 1 |
| 10 | 73494096 | G | A | p.Val1402Met | CDH23 | T | D | P | D | D | M | . | T | T | 0.201 | 22.2 | 4.49 | 1 |
| 10 | 73571298 | T | C | p.Phe3077Leu | CDH23 | D | B | B | D | D | M | T | T | T | 0.157 | 15.52 | 3.05 | 1 |
| 10 | 73571717 | C | T | p.Arg3109Cys | CDH23 | D | D | B | D | D | L | T | T | T | 0.22 | 17.51 | 5.82 | 1 |
| 10 | 73468881 | G | C | p.Val1045Leu | CDH23 | D | B | B | D | D | L | . | T | T | 0.249 | 19.95 | 2.74 | 1 |
| 10 | 73330674 | C | T | p.Pro251Leu | CDH23 | T | B | B | D | D | L | T | T | T | 0.051 | 13.43 | 4.28 | 1 |
| 10 | 73498394 | G | A | p.Ser1450Asn | CDH23 | T | B | B | D | D | N | . | T | T | 0.031 | 11.65 | 5.38 | 1 |
| 10 | 73544798 | C | T | p.Arg1885Cys | CDH23 | T | P | P | N | D | M | . | T | T | 0.355 | 17.41 | 3.5 | 1 |
| 10 | 73553202 | G | A | p.Glu2173Lys | CDH23 | T | P | B | D | D | N | . | T | T | 0.196 | 21 | 5.28 | 1 |
| 10 | 73567077 | C | T | p.Ser2741Leu | CDH23 | T | B | P | D | D | L | T | T | T | 0.16 | 26.5 | 4.77 | 1 |
| 10 | 73337686 | A | G | p.Ile257Val | CDH23 | T | B | B | N | D | N | T | T | T | 0.047 | 9.05 | -2.29 | 1 |
| 10 | 73439175 | A | T | p.Gln595Leu | CDH23 | T | B | B | N | D | N | . | T | T | 0.063 | 13.46 | 3.78 | 1 |
| 10 | 73491830 | G | A | p.Val1268Met | CDH23 | T | B | B | N | D | L | . | T | T | 0.121 | 11.74 | 3.9 | 1 |
| 10 | 73500622 | A | G | p.His1511Arg | CDH23 | T | B | B | N | D | N | . | T | T | 0.201 | 13.23 | 5.06 | 1 |
| 10 | 73468899 | G | A | p.Val1051Ile | CDH23 | T | B | B | N | N | N | . | T | T | 0.044 | 5.205 | 1.04 | 1 |
| 10 | 73550915 | A | G | p.Ile2026Val | CDH23 | T | B | B | N | N | N | . | T | T | 0.037 | 2.379 | -6.59 | 1 |

Annotations: SIFT, Polyphen2_HDIV, Polyphen2_HVAR, LRT, MutationTaster, MutationAssessor, FATHMM, RadialSVM, LR
https://annovar.openbioinformatics.org/en/latest/user-guide/filter/#ljb42-dbnsfp-non-synonymous-variants-annotation

| | |
|---|---|
| SIFT (sift) | D: Deleterious (sift<=0.05); T: tolerated (sift>0.05) |
| PolyPhen 2 HDIV (pp2_hdiv) | D: Probably damaging (>=0.957), P: possibly damaging (0.453<=pp2_hdiv<=0.956); B: benign (pp2_hdiv<=0.452) |
| PolyPhen 2 HVar (pp2_hvar) | D: Probably damaging (>=0.909), P: possibly damaging (0.447<=pp2_hdiv<=0.909); B: benign (pp2_hdiv<=0.446) |
| LRT (lrt) | D: Deleterious; N: Neutral; U: Unknown |
| MutationTaster (mt) | A" ("disease_causing_automatic"); "D" ("disease_causing"); "N" ("polymorphism"); "P" ("polymorphism_automatic") |
| MutationAssessor (ma) | H: high; M: medium; L: low; N: neutral. H/M means functional and L/N means non-functional |
| FATHMM (fathmm) | D: Deleterious; T: Tolerated |
| RadialSVM | D: Deleterious; T: Tolerated |
| LR | D: Deleterious; T: Tolerated |
| REVEL | higher scores are more deleterious |
| CADD_phred | higher scores are more deleterious |
| GERP++ (gerp++) | higher scores are more deleterious |

**Supplementary Figure 9: *CDH23* variants Identified in Japanese individuals with schizophrenia**. Patient-specific novel rare loss of function (LoF) variants predicted by *in silico* tools. There variants were very rare and observed in individual cases.

A

B

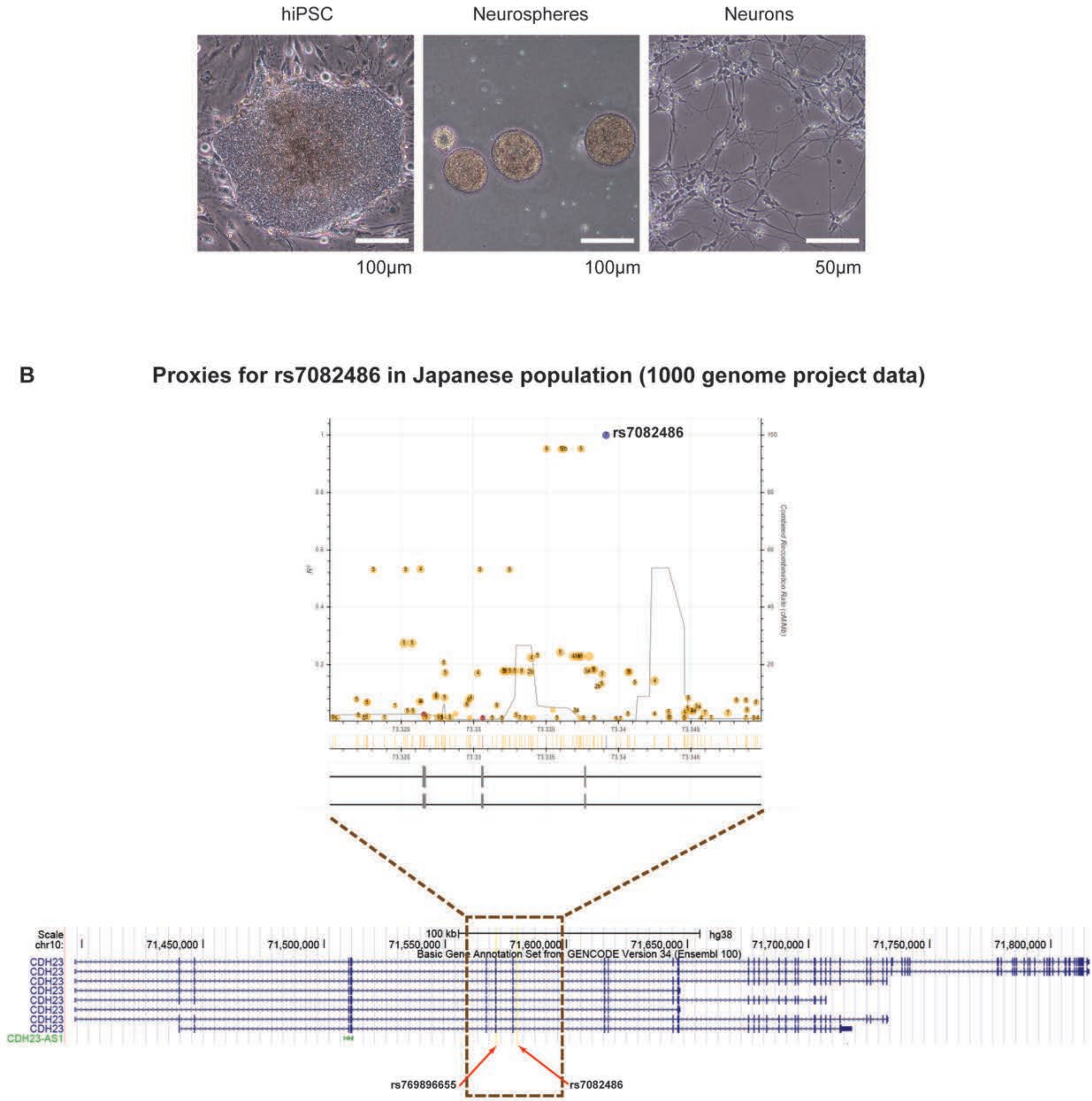


**Supplementary Figure 10: hiPSCs generated from a healthy individual with rs769896655 (c.753G>A) variant and the linkage disequilibrium pattern of the PPI GWAS hit in Japanese population (A)** Bright field images of hiPSC established from a healthy subject (TKUR120) heterozygous (G/A) for the variant rs769896655, neurospheres, and neurons derived from the hiPSC are shown, which was used for testing the allele specific expression in *CDH23* transcript. (**B**) Significant variant from the GWAS of PPI in schizophrenia, rs7082486 in *CDH23,* was located downstream of the the variant rs769896655 (c.753G>A) with considerable linkage disequilibrium (LD) in Japanese population (https://ldlink.nci.nih.gov/?tab=home). Since the variant rs769896655 was not represented in the 1000genome data, the exact LD value could not be estimated

**List of supplementary tables**

**Supplementary table 1:** Strain-specific single nucleotide variants located on the previously reported loci for PPI, selected for genotyping

**Supplementary Table 2:** Sequence for short guide RNA (sgRNA) and single-stranded-oligodeoxynucleotide (ssODN) for generating CRISPR/Cas9n mediated knock-in mouse

**Supplementary Table 3:** Primer sequences for the gene expression analysis

**Supplementary Table 4:** Molecular inversion probes sequences targeting the coding exons and the flanking exon-intron boundaries of *CDH23*

**Supplementary Table 5:** QTL mapping performed for 86 dB PPI by composite interval mapping with LOD score and percentage of variance explained for the phenotype (R2) in 1,010 F2 mice derived from F1 parents (B6NjC3Hj), both combined and dichotomized for gender

**Supplementary Table 6:** Bayesian multiple QTL mapping performed for 86 dB-PPI with posterior probabilities across additive and dominant models along with percentage of variance explained (R2) in 1,010 F2 mice derived from F1 parents (B6NjC3Hj), both combined and dichotomized for gender

**Supplementary Table 7**: Single nucleotide variants, which were distinct between the C57BL/6NJ and C3H/HeJ mouse strains, within the chromosome 10 QTL region from the sequence variation database of mouse strains (https://www.sanger.ac.uk/science/data/mouse-genomes-project/; REL-1303- GRCm38)

**Supplementary Table 8:** Variant allele frequency of the *CDH23* rs769896655 among different populations

**Supplementary Table 9:** Association analysis of *CDH23* variant rs769896655 with schizophrenia (pooled samples)
